# Single-cell transcriptomics of the aged mouse brain reveals convergent, divergent and unique aging signatures

**DOI:** 10.1101/440032

**Authors:** Methodios Ximerakis, Scott L. Lipnick, Sean K. Simmons, Xian Adiconis, Brendan T. Innes, Danielle Dionne, Lan Nguyen, Brittany A. Mayweather, Ceren Ozek, Zachary Niziolek, Vincent L. Butty, Ruth Isserlin, Sean M. Buchanan, Stuart R. Levine, Aviv Regev, Gary D Bader, Joshua Z. Levin, Lee L. Rubin

## Abstract

The mammalian brain is complex, with multiple cell types performing a variety of diverse functions, but exactly how the brain is affected with aging remains largely unknown. Here we performed a single-cell transcriptomic analysis of young and old mouse brains. We provide a comprehensive dataset of aging-related genes, pathways and ligand-receptor interactions in nearly all brain cell types. Our analysis identified gene signatures that vary in a coordinated manner across cell types and gene sets that are regulated in a cell type specific manner, even at times in opposite directions. Thus, our data reveals that aging, rather than inducing a universal program drives a distinct transcriptional course in each cell population. These data provide an important resource for the aging community and highlight key molecular processes, including ribosomal biogenesis, underlying aging. We believe that this large-scale dataset, which is publicly accessible online (aging-mouse-brain), will facilitate additional discoveries directed towards understanding and modifying the aging process.

## INTRODUCTION

Aging, the time-dependent functional decline of organs and tissues, is the biggest risk factor for many diseases, including several neurodegenerative and cardiovascular disorders ^1^. Characterizing aging-related molecular and cellular changes will provide insights into this complex process and highlight opportunities to slow or reverse its progression, thereby helping to prevent or treat aging-associated pathologies. That this might be achievable is supported by a plethora of studies using model organisms demonstrating that not only lifespan, but also the integrity of multiple tissues, can be regulated by discrete molecular modifications ^2–4^.

Towards the goal of achieving a broader understanding of aging-related changes and deciphering the molecular mechanisms that accompany brain aging, transcriptomic studies in model organisms and humans have been at the forefront of experiments. However, these studies generally utilize aggregated RNA from either mixed cell populations ^5–10^ that may vary in distinct ways with age, or cell populations purified using known markers ^11–15^, which themselves may also change during aging. Therefore, despite the successful identification of major aging-related genes and pathways, prior transcriptomic analyses have not resolved the common aging-related changes experienced across all brain cells from those that may be cell-type specific. Thus, there is a need to elucidate how individual cell types are affected by aging and to clarify if the process of aging follows a similar blueprint in all cell types or whether certain cell types have unique transcriptional changes. This will be critical in determining whether aging at the tissue level is a global process, if it results from specific changes in certain cell types that culminate in loss of function and deterioration, or a combination of both ^16^. This information may also help the design of effective aging-related therapeutics that are targeted either narrowly - affecting only certain types of cells - from those that are targeted more broadly.

In this study, to begin to address these issues, we employed single-cell RNA sequencing to profile and compare the cellular composition and transcriptomes of young and old mouse brains. This is the first large-scale transcriptomic analysis of aging for the vast majority of individual brain cell types. For all the major cell populations identified, we provide a comprehensive dataset of genes and pathways whose transcriptional profiles change with aging. Our computational analysis suggests that cells in the brain do not change with aging identically, indicating that, while overlapping signatures exist, the cellular consequences of aging are not universal. Given that cell non-autonomous changes are also known to regulate aging-dependent changes ^2,3^, we also detail ligand-receptor interactions among nearly all the cell types in the brain that are modified by aging. Overall, this study provides a rich resource that can facilitate the interrogation of the molecular underpinnings and cellular basis of the aging process in the mouse brain.

## RESULTS

### Identification of cell types

To gain new, more precise, insights into the effects of aging, we employed unbiased high-throughput single-cell RNA sequencing (scRNA-seq) to examine the transcriptional profiles of young and old mouse brains (Fig. 1A). Because the dissociation of mammalian adult brains is challenging due to the complexity of the tissue involving a very dense intercellular space, including an extensive neuronal network insulated by myelin sheaths, we first developed a new dissociation protocol that enables the isolation of healthy and intact cell suspensions that are representative of both young and old mouse brains (see details in Supplementary Methods).

**Fig. 1.**
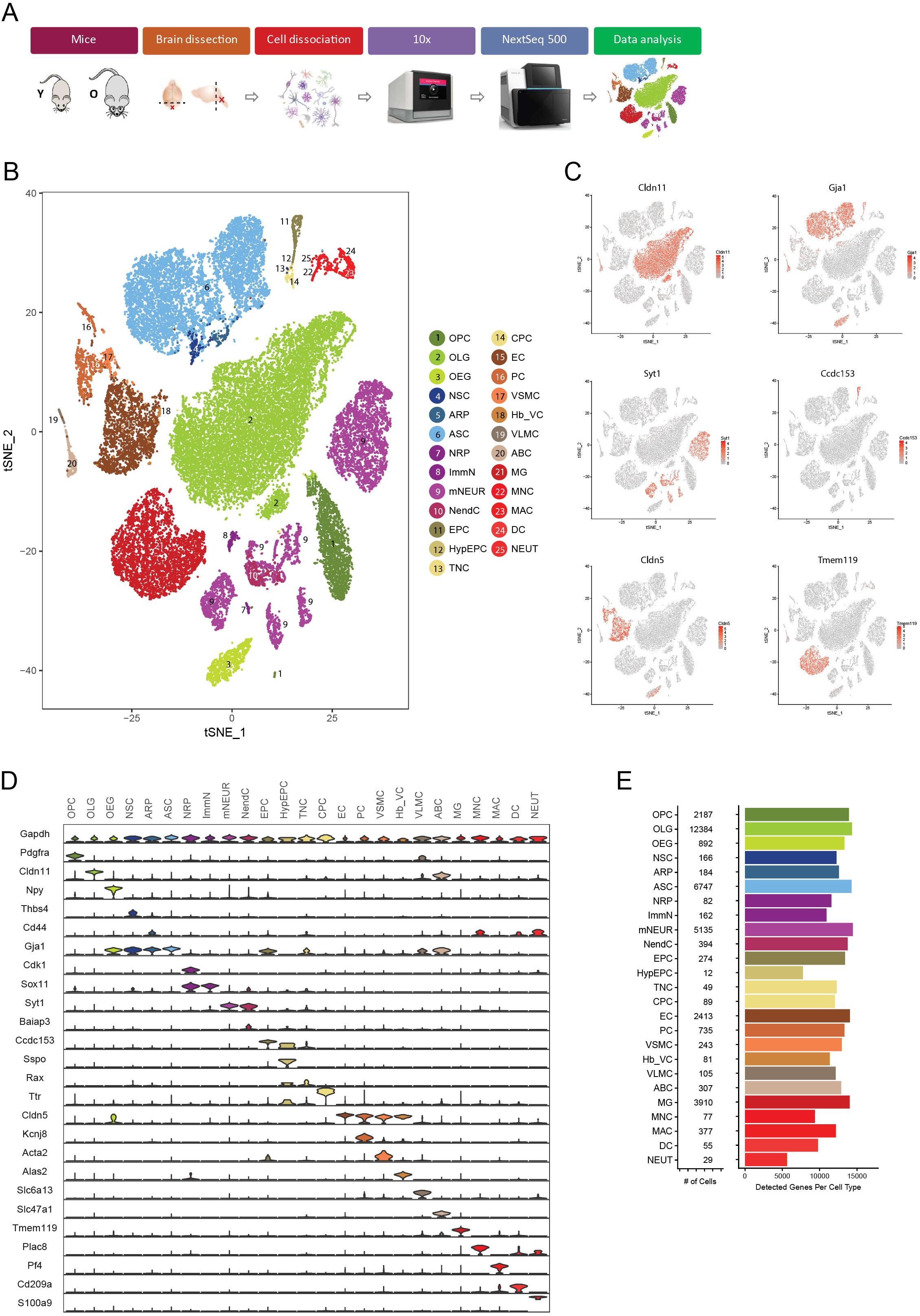
Identification of cell types. (A) Overview of the experimental workflow. (B) t-distributed stochastic neighbor embedding (t-SNE) projection of 37,089 cells derived from 8 young and 8 old mouse brains. Cell clusters were color-coded and annotated post hoc based on their transcriptional profile identities (see Supplementary Methods). (C) t-SNE visualization of 6 major cell populations showing the expression of representative well-known cell-type-specific marker genes. Numbers reflect the number of unique molecular identifiers (UMI) detected for the specified gene for each cell. (D) Violin plot showing the distribution of expression levels of known representative cell-type-enriched marker genes across all 25 cell types. Markers used: *Pdgfra*, OPC ^22^; *Cldn11*, OLG ^22^; *Npy*, OEG ^29^; *Thbs4*, NSC ^72^–^75^; *Cd44*, ARP ^76^; *Gja1*, ASC ^26^; *Cdk1*, NRP ^28^; *Sox11*, ImmN ^28^; *Syt1*, mNEUR ^21^; *Baiap3*, NendC ^77^; *Ccdc153*, EPC ^23^; *Sspo*, HypEPC ^32^; *Rax*, TNC ^23^; *Ttr*, CPC ^19^; *Cldn5*, EC ^30^; *Kcnj8*, PC ^30^; *Acta2*, VSMC ^30^; *Alas2*, Hb-VC ^27,78^; *Slc6a13*, VLMC ^22,32^; *Slc47a1*, ABC ^32^; *Tmem119*, MG ^26^; *Plac8*, MNC ^31^; *Pf4*, MAC ^31^; *Cd209a*, DC ^31^,^79^; *S100a9*, NEUT ^31^. (E) Bar plot showing the total number of detected cells and the total number of detected genes per cell type.

We then analyzed the transcriptomes of 50,212 single cells (24,401 young and 25,811 old) derived from the brains of 8 young (2-3 months) and 8 old (21-23 months) mice (Fig. S1-2). In our initial analysis, we leveraged a graph-based clustering approach ^17,18^ to aggregate transcriptionally similar cells. Next, we removed clusters likely to be of low quality, resulting from debris, doublets/multiplets and dead cells (Fig. S3), and employed other critical quality control steps as described in the Supplementary Methods (Fig. S4). Ultimately, our analysis led to the identification of 37,089 cells (Fig. S5A), representing 25 cell populations (Fig. 1B) with distinct expression profiles (Fig. 1C-D and Fig. S6): oligodendrocyte precursor cells (OPC), oligodendrocytes (OLG), olfactory ensheathing glia (OEG), neural stem cells (NSC), astrocyte-restricted precursors (ARP), astrocytes (ASC), neuronal-restricted precursors (NRP), immature neurons (ImmN), mature neurons (mNEUR), neuroendocrine cells (NendC), ependymocytes (EPC), hypependymal cells (HypEPC), tanycytes (TNC), choroid plexus epithelial cells (CPC), endothelial cells (EC), pericytes (PC), vascular smooth muscle cells (VSMC), hemoglobin-expressing vascular cells (Hb-VC), vascular and leptomeningeal cells (VLMC), arachnoid barrier cells (ABC), microglia (MG), monocytes (MNC), macrophages (MAC), dendritic cells (DC), and neutrophils (NEUT). Cell counts and other sequencing metrics for each identified cell type are shown in Fig. 1E and Fig. S5B-E.

### Identification of cell subtypes and states

In order to reveal heterogeneity within each population, we grouped the aforementioned cell types based on their expression profile, lineage, function and anatomical organization into 6 classes (oligodendrocyte lineage, astrocyte lineage and stem cells, neuronal lineage, ependymal cells, vasculature cells, and immune cells) (Fig. S7) and employed another round of clustering. This subsetting of the dataset enabled us to highlight more subtle changes within the classes without the impact of variation due to inclusion of drastically different cell identities. This secondary analysis identified dozens of different cell subtypes and states reflecting distinct functional, maturational and regional cell identities (Fig. S8-9). All of these cell identities are in line with recent scRNA-seq studies ^19–33^, whose purpose was to identify novel and distinct cell types/subtypes and create detailed atlases of the developing and adult mouse brain (see details Fig. S8). This allowed us to generate a comprehensive dataset of gene expression profiles for all the experimentally validated cell types from both young and old brains (Table S1 and 2) at a resolution comparable to that of the aforementioned studies. This also permitted us to identify specific markers that distinguish each type regardless of age (Table S3).

### Aging-related effects on cell-to-cell transcriptional variability and cellular composition

We found that cell identity is largely preserved in young and old brains as indicated by unbiased clustering where all clusters represent cells of all animals from both ages (Fig. S4C). Furthermore, the quality of data generated from both young and old cell types appear similar, with each having comparable numbers of unique molecular identifiers (UMIs) and detected genes (Fig. S5C, E). Next, we compared the coefficient of variation (CV) of expression for all of the transcribed genes (Fig. S10A), only the mitochondrially-encoded genes (Fig. S10B), or only the ribosomal genes (Fig. S10C). We observed differences in the variability of transcription between young and old cells in many cell types. However, the directionality of change was not consistent among cell types, providing evidence that aging is not broadly associated with increased transcriptional variation ^34^.

Then, by investigating the abundance of each cell type, we found that cellular composition was largely consistent across both young and old brains (Fig. 2A and Table S4). Nonetheless, we were able to confirm the previously reported aging-related decline of certain cell populations, such as NRP ^35,36^ (Fig. 2A), and to reveal potentially interesting but not statistically significant population shifts within certain subtypes of OPC, OLG, ASC and MG (Fig. S11, see also Fig. S8B, F). Of note, although the estimated percentages for each cell type do not necessarily reflect their actual proportions in the mouse brain, mainly due to differences in their sensitivity to tissue dissociation, the observed changes in cell type ratios appear to reflect a real biological effect.

**Fig. 2.**
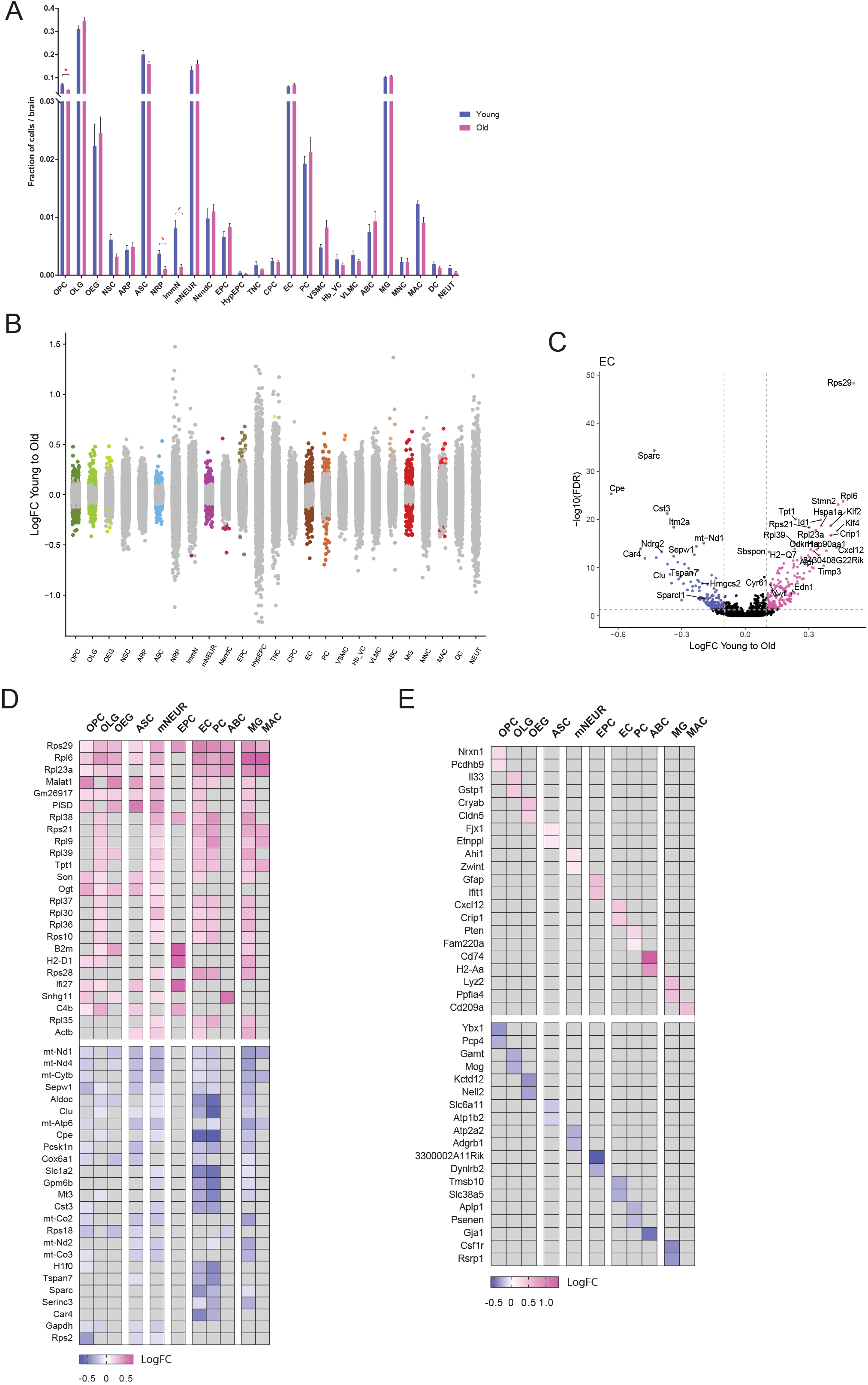
Aging-related population shifts and changes in gene expression. (A) Bar plot showing the fraction of cells associated with each cell type in both young and old brains (data presents mean ± SEM of 8 young and 8 old brains; *FDR<0.05 by Mann-Whitney *U* test). (B) Strip chart showing the aging-related logarithmic fold changes (logFC) of all detected genes across all cell types. Each point represents a gene. Genes in colored dots are significantly (FDR<0.05) upregulated (logFC>0.1) or downregulated (logFC<−0.1) with aging. Genes in gray are not significantly changed with aging. (C) Sample volcano plot for EC showing-log10(FDR) and logFC values for all genes with highlighting for those that are significantly upregulated (logFC>0.1; magenta dots) or downregulated (logFC<−0.1; blue dots) with aging. Genes in black are not significantly changed with aging. (D) Heatmap of logFC showing a subset of aging-related genes (FDR<0.05 and FC>10%) that are shared across many of the major cell types. Gray indicates no significant dysregulation. (E) Heatmap of logFC showing a subset of aging-related genes (FDR<0.05 and FC>10%) that are unique to each major cell type.

### Identification of aging-related genes

We then investigated the breadth of transcriptional changes that occur in the mouse brain with aging by performing differential gene expression (DGE) analysis between young and old cell types for all 25 distinct cell populations (Table S5). Of the 14,699 total detected genes, 3,897 were significantly affected by aging in at least one cell type (FDR<0.05). When the magnitude of change in expression was also considered, 1,117 genes passed the 10% fold-change (FC) threshold (Fig. 2B and Table S6). Interestingly, of those, 1,034 exhibited the same directionality regardless of the cell type identity (494 upregulated and 540 downregulated), while the direction of change in the expression of 83 genes was different across cell populations (discussed further below; Table S6). As described in Supplementary Methods, our ability to identify genes whose transcription changes significantly with aging and the calculation of fold-change is dependent on several factors, including the number of cells within each population, the level of transcription, and the algorithm for analysis.

### Identification of shared and cell-type specific aging signatures

To explore the validity of these aging signatures, we first started broadly and compared our data with past transcriptomic studies of the mouse aging brain ^5,7,37^. To more effectively compare datasets, we aggregated all our sequenced cells recreating a traditional whole brain profile similar to what might be observed using bulk sequencing (Table S2 and 5). As expected, this analysis verified previously identified top aging-related genes (such as *Apod*, *B2m*, *C1qa*, *C4b*, *Ctss*, *Il33*, *Rpl8*). Moreover, due to the increased sensitivity of the techniques used in our study compared to past ones, we were able to identify a set of novel aging-related genes not reported previously (such as *Apoc1*, *Caly*, *Cxcl12*, *Nell2*, *Ybx1;* presented in Table S5). These changes could have been masked in past studies due to their limited expression levels or variations in less abundant cell types. Importantly, our single cell DGE data enabled us to build on these results to identify from which cell types these aging signatures arose. For example, *Ctss*, while highly transcribed in all immune cells (MG, MAC, MNC, DC; see Table S2), was only significantly changed with aging in MG (Table S5). Another example is *Nell2*, which is mostly transcribed in neuronal lineage cells and OEG, but its levels changed with aging only in OEG.

We then focused our analysis on 11 major cell populations that exhibited the greatest number of differentially regulated genes (Fig. 2B). By comparing the DGE data from these cell populations (Fig. 2C and Fig. S12), we were able to distinguish both shared and cell-type specific aging signatures. Table S6 presents a matrix that specifies the genes that changed significantly in each cell type.

Fig. 2D presents selected top aging-related genes that are shared across multiple cell types. The majority of the most commonly aging-upregulated genes were ribosomal genes (such as *Rpl6*), lncRNA genes(such as *Malat1*)and immunoregulatory/inflammatory genes (such as *B2m*). The most commonly aging-downregulated genes were mitochondrial respiratory chain complex genes (such as *mt-Nd1*), glycolysis-related genes (such as *Aldoc*), genes encoding selenoproteins (such as *Sepw1*) and tetraspanins (such as *Tspan7*) (Fig. 2D and Table S6). As shown in Fig. 3A, we were able to validate the transcriptional changes of representative commonly aging-related genes by employing in situ hybridization assays in mouse brain sections.

**Fig. 3.**
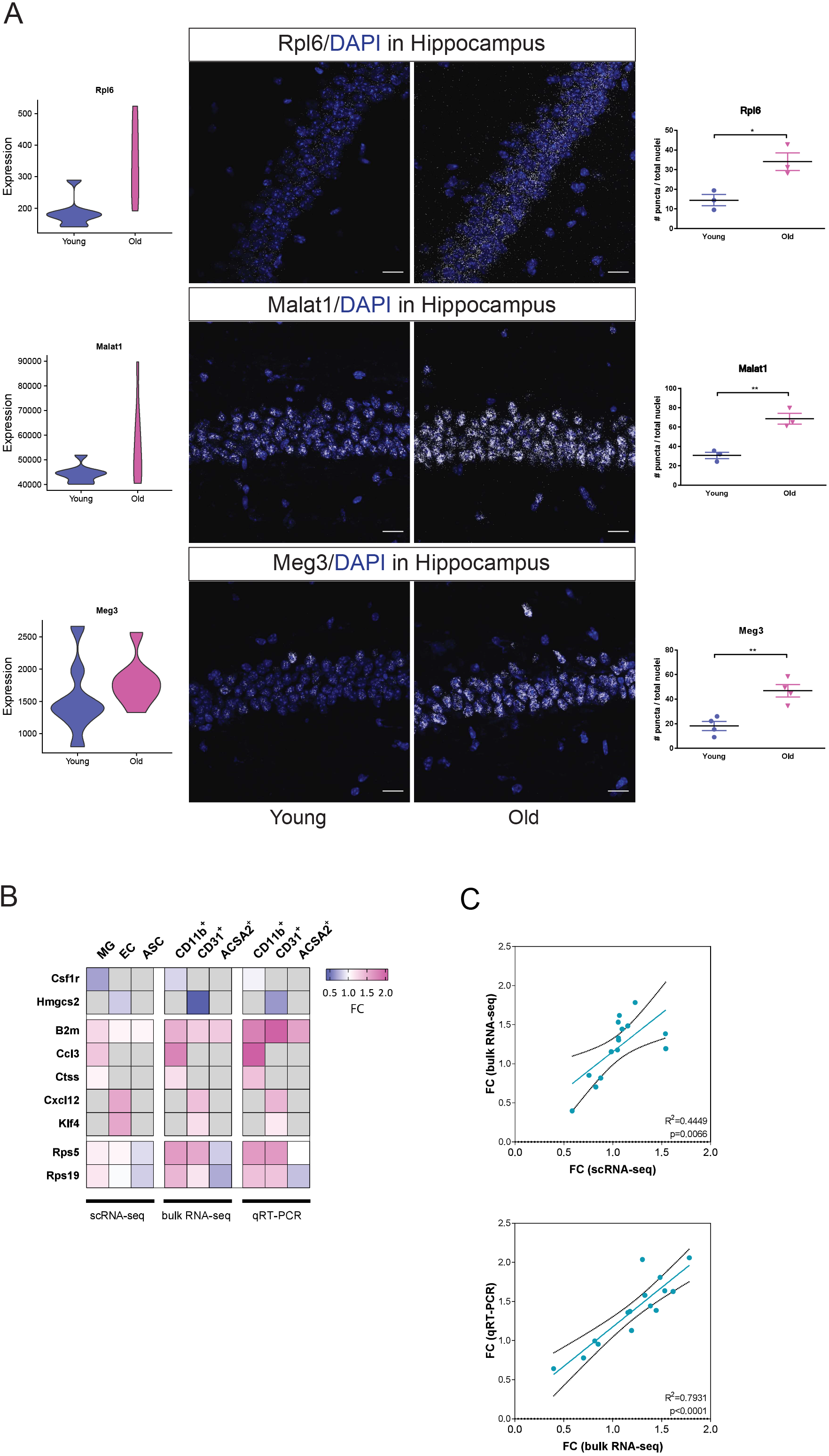
Validation of representative aging-related gene expression changes. (A) Violin plots with data in units of transcripts per million (TPM) from our scRNA-seq across all identified cell types (left panels) and RNAscope in situ hybridization images (middle panels) of mouse hippocampi showing the aging-related upregulation of *Rpl6*, and of the lncRNAs *Malat1* and *Meg3*. This brain area was selected due to its high expression levels of those genes, according to the Allen Brain Atlas ^80^. Scatter plots (right panels) showing the quantification of the RNAscope data of 3-4 independent experiments (data presents mean ± SEM; *p<0.05, **p<0.01 by Welch’s t-test). Scale bar: 20um. (B) Heatmap showing the fold expression changes (FC) of a few representative significantly (FDR<0.05) aging-related genes in MG, EC, and ASC as identified by our scRNA-seq (left panel), and verified by both bulk RNA-seq (middle panel) and qRT-PCR (right panel) on sorted CD11b^+^, CD31^+^ and ACSA-2^+^ cells. Gray indicates no or minimal expression in the seq data and no amplification in the qRT-PCR data. For the qRT-PCR experiments data presents mean ± SEM of 3-5 young and 3-6 old mouse brains. (C) Scatter plots showing the significant correlations of the gene expression changes in (B) between the scRNA-seq, bulk RNA-seq and qRT-PCR datasets. (D) Violin plots and immunohistochemistry showing the aging-related up-regulation of IL33 (mainly expressed in OLG based on our scRNA-seq and others ^24,27^), and the aging-related down-regulation of SPARC in MG (IBA1-positive cells; indicated by arrows). Scale bar: 50um. Scatter plots showing the quantification of the immunohistochemistry data of 4 independent experiments (data presents mean ± SEM; *p<0.05 by Welch’s t-test).

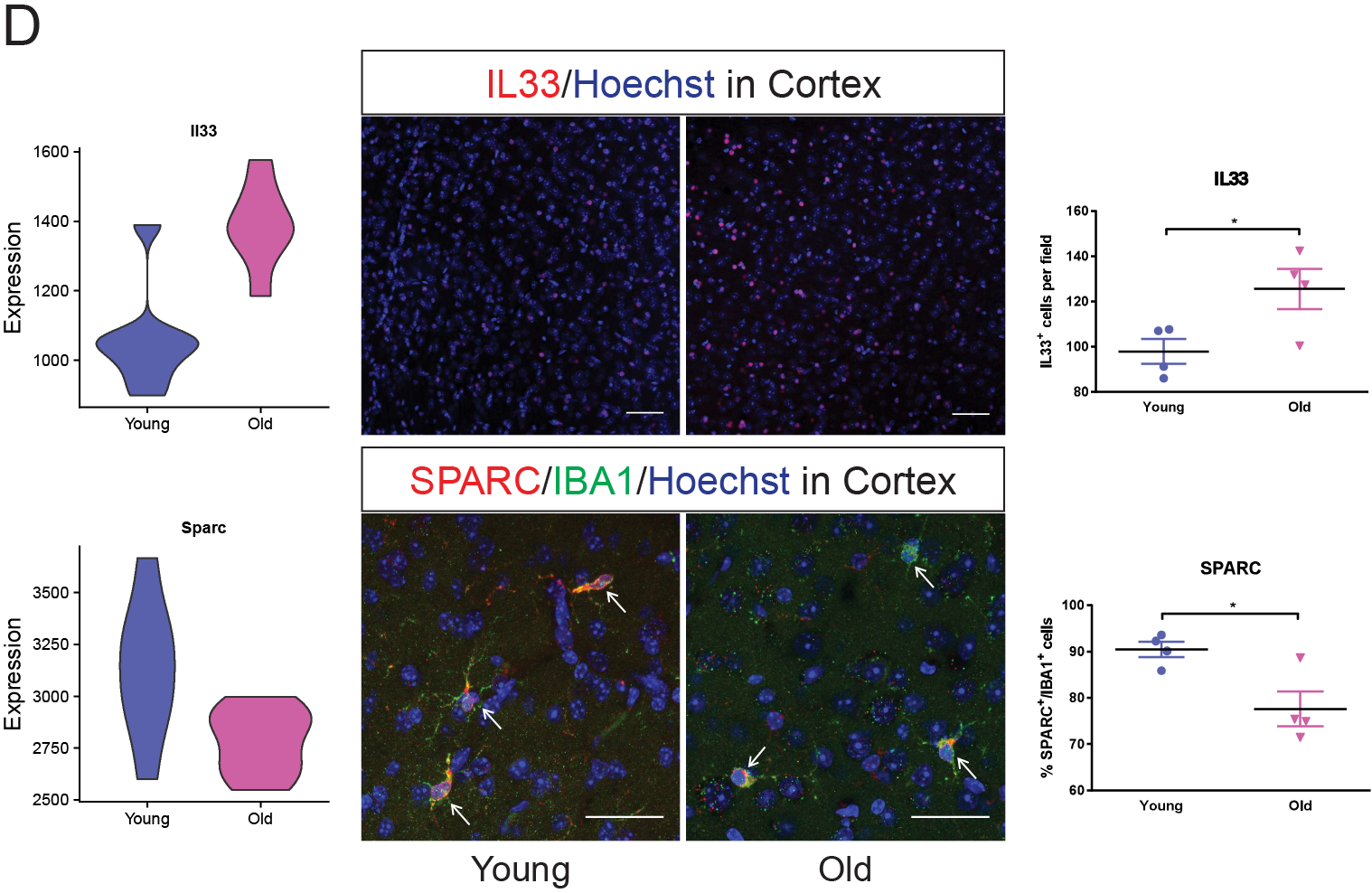

A fraction of genes representing cell-type specific aging signatures are highlighted in Fig. 2E. Interestingly, these data revealed that certain genes that are traditionally used as cell type specific markers change with aging, such as the aging-related downregulation of *Mog* in OLG and *Csf1r* in MG, and the aging-related upregulation of *Cxcl12* in EC (Fig. 2E and Table S5). Conversely, we observed that other classic cell type markers that are used to specify certain populations change with aging in others. For example, *Gfap*, which is highly transcribed and enriched in the astrocyte lineage and stem cells (Table S2 and 3), was found as one of the top aging-related genes in EPC (Fig. 2E and Table S5).

To validate certain common and cell type unique aging signatures, we employed both bulk RNA-seq and qRT-PCR analysis on FACS-purified EC, MG and ASC from both young and old brains (Fig. S13). As shown in Fig. 3B-C, we verified the aging-related changes of specific genes (such as *B2m*, *Csf1r*, *Ctss*, *Cxcl12*) in their corresponding cell types. Additionally, to further assess whether our transcriptomic approach faithfully captured changes at the protein level we performed immunohistochemistry assays. As shown in Fig. 3D, we verified the aging-related downregulation of SPARC in MG and the broad aging-related upregulation of IL33, whose expression was shown to change in the whole brain data with the majority of signal arising from OLG, as revealed by the single cell data (Fig. 2E and Table S2 and 5).

### Identification of bidirectional aging signatures

Interestingly, data analysis revealed genes with opposite regulation between different cell types. An example is *Cldn5* that is often used as a marker for vasculature cells, specifically for EC, but it is also highly transcribed in OEG (Table S2). We found aging-related downregulation of *Cldn5* in EC but upregulation in OEG. Notably, when its levels were measured broadly in the whole brain, changes were minimal (Table S2 and 5). This is important as it explains why certain aging-related genes were masked in prior bulk sequencing studies.

Similarly, we found large gene sets that were discordant among groups of cell types. One of these referred to ribosomal genes. Many ribosomal genes were found among the top most common aging-related ones across major cell populations (Fig. 2D and Table S6), but a subset of these genes was also identified as genes that exhibited differential regulation/directionality with aging in a cell-type specific manner (Fig. S14 and Table S6; see also Fig. 3B). When we examined the expression profile of all genes encoding ribosomal proteins among major cell populations we found two distinct and divergent patterns. As shown in Fig. 4A, both OPC and ASC were found to downregulate a fraction of their ribosomal genes with aging, while the other cell types upregulated their expression. To validate these bidirectional aging-related changes, we examined ribosomal gene expression in FACS-purified ASC, EC and MG. As shown in Fig. 4B-C, bulk RNA-seq recapitulated the scRNA-seq data for a subset of key genes, thus verifying the existence of divergent aging-driven ribosomal expression patterns among the tested cell populations and highlighting their potentially distinct responses to aging.

**Fig. 4.**
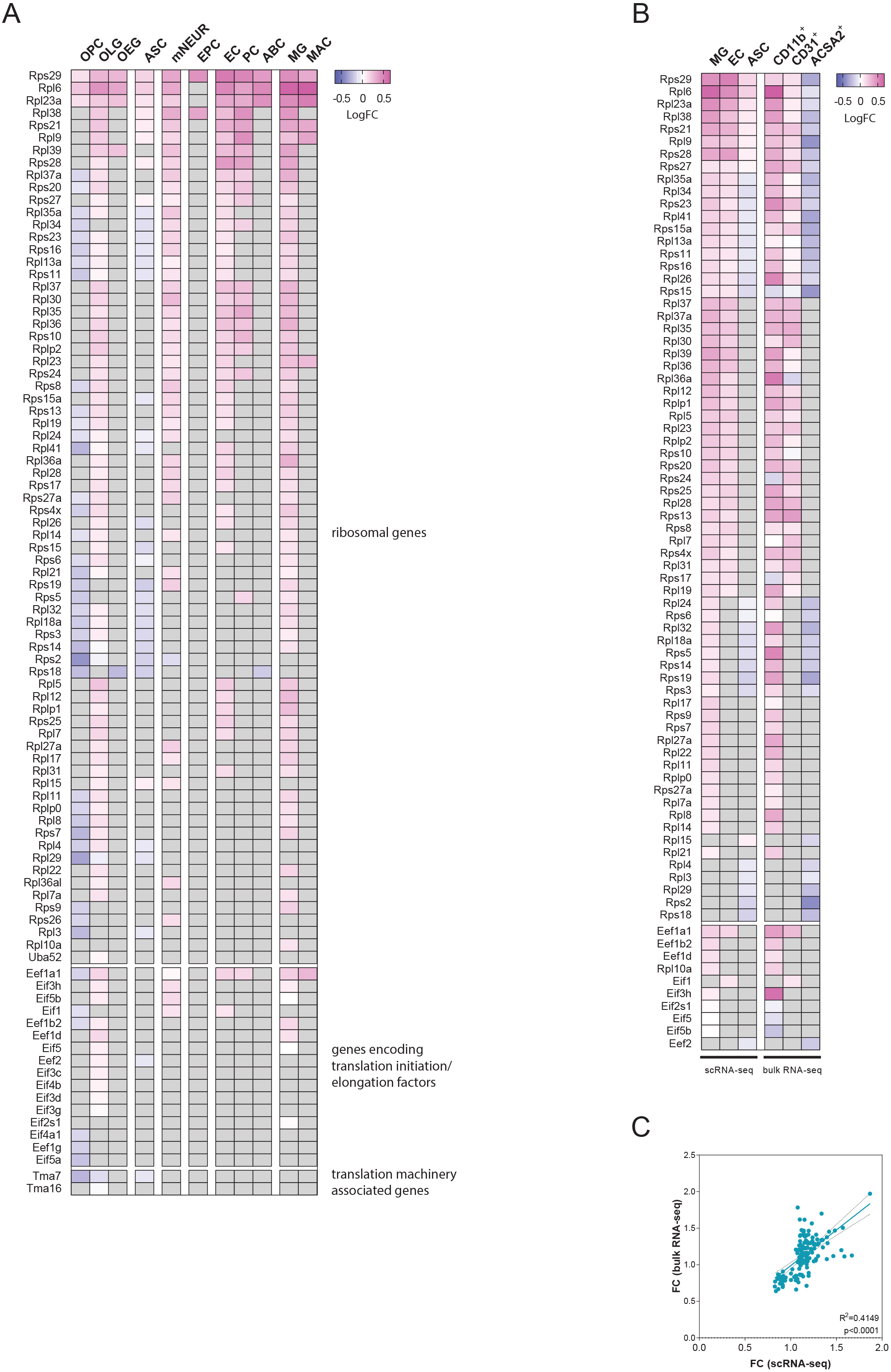
Aging-related bidirectional changes in the expression of ribosomal genes. (A) Heatmap showing the logFC for all the significantly (FDR<0.05) aging-related ribosomal and translation-associated genes across 11 cell types, as identified by our scRNA-seq. (B) Heatmap of logFC showing all the significant (FDR<0.05) aging-related ribosomal and translation-association genes across MG, EC and ASC as identified by our scRNA-seq (left panel), and further verified by bulk RNA-seq (right panel) on sorted CD11b^+^, CD31^+^ and ACSA-2^+^ cells. Despite the fact that only a subset of these genes was found significantly dysregulated in our bulk RNA-seq analysis, due to lower statistical power, there is a significant correlation of the gene expression changes between the scRNA-seq and bulk RNA-seq datasets, as shown in scatter plot (C). More specifically, dots in (C) represent all genes from the examined cell types in (B). Linear regression is depicted with the colored line, while black dotted lines represent 95% confidence intervals. Correlation coefficient (R^2^) and p-value are shown at the bottom right part of the plot.

### Identification of aging-related pathways

Next, we investigated changes in aging-related cellular pathways and processes by performing gene set enrichment analysis (GSEA) ^38^. GSEA has increased sensitivity compared to DGE analysis as it aggregates information from broad sets of genes that are presumed to be functionally related. As such, we were also able to include cell types with limited cell numbers that did not show significant aging-related changes by DGE analysis. This approach revealed the existence of many shared and cell-type specific aging-related pathways across the examined cell populations (Fig. 5 and Table S7 and 8). In total, 451 pathways (1,142 GSEA terms) changed significantly (p<0.05 & q<0.25); 234 were expressed in at least 2 cell types, while the remaining 217 were unique for specific cell populations. Of the 451 aging-related pathways, 339 exhibited the same directionality regardless of cell type identity (195 were upregulated and 144 downregulated), while the directionality of changes in the remaining 112 varied across cell types (Table S8). The most common aging-related pathways were those associated with cellular respiration, protein synthesis, inflammatory response, oxidative stress, growth factor signaling, and neurotrophic support (Fig. 5 and Table S8).

**Fig. 5.**
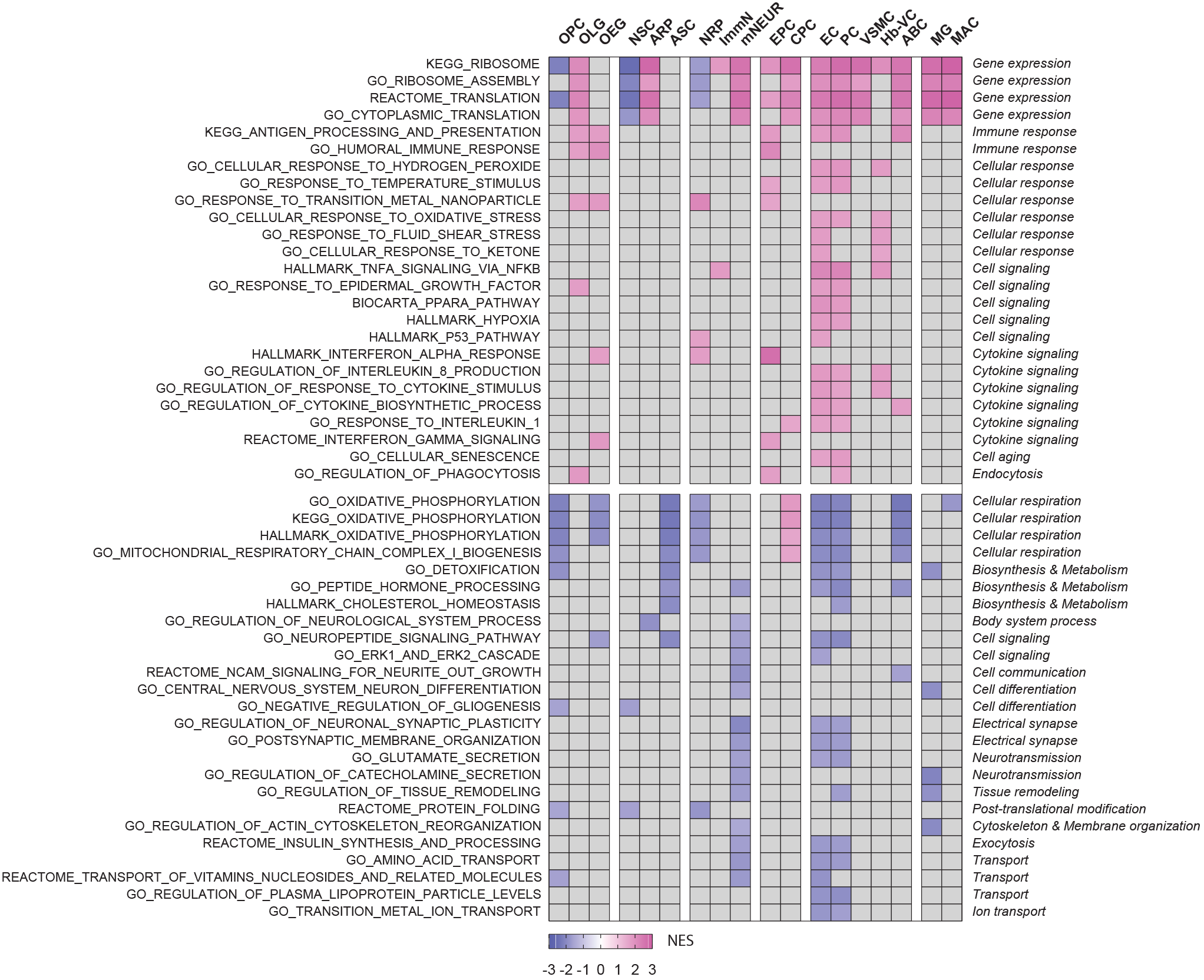
Aging-related changes in cellular pathways and processes. Heatmap of gene set enrichment analysis (GSEA) showing a small subset of significant (p<0.05 and q<0.25) aging-related pathways across major cell types. Numbers in legend correspond to normalized enrichment scores (NES). Positive NES values indicate upregulation, while negative NES values indicate downregulation. Gray indicates no significant dysregulation.

Here, we highlight changes in EC and EPC, two understudied, but important, brain cell populations, due to their unique roles in forming the barriers that isolate the brain parenchyma from factors circulating in blood and cerebrospinal fluid. GSEA showed that EC exhibit numerous aging-regulated cellular pathways and biological processes, such as the aging-related induction of senescence, hypoxia signaling and response to ketone signaling, and the aging-related reduction of xenobiotic metabolism, lipid metabolism and hormone processing (Fig. S15A and Table S7 and 8). In the EPC population, a notable finding in the pathway analysis is the aging-related upregulation of interferon signaling (Fig. S15B and Table S7 and 8). This aligns with the induction of certain interferon-stimulated genes (like *Ifit1*) that was shown in the DGE analysis (Fig. 2E and Table S5) and is similar to what has been previously reported for the choroid plexus epithelium ^39^. Thus, this unique aging signature seems to define transcriptional specialization in EPC and may point to their participation in aging-related inflammatory responses in the brain parenchyma.

Importantly, in line with our DGE analysis, GSEA highlights that ribosomal biogenesis is a biological process that exhibits differential regulation with aging across different cell populations, beyond what we found based on DGE analysis alone. In particular, this pathway analysis, despite employing stringent significance criteria, clearly showed that, with aging, the vast majority of the differentiated brain cell types exhibited upregulation of genes encoding ribosomal subunits, while the majority of progenitor cells exhibited the opposite regulation (Fig. 5, Fig. S16 and Table S8).

### Identification of aging-related changes in intercellular communication

Lastly, our single-cell transcriptomics data provides the ability to explore how aging-driven changes in gene expression could affect intercellular communication within the brain. By leveraging the transcriptional profiles of each cell type, we built a comprehensive intercellular network of potential ligand-receptor interactions among nearly all the identified brain cell types (Fig. 6A). We then enriched this network with data from our DGE analysis to mark all those interactions that were found to change with aging at the ligand or receptor level (Table S9).

**Fig. 6.**
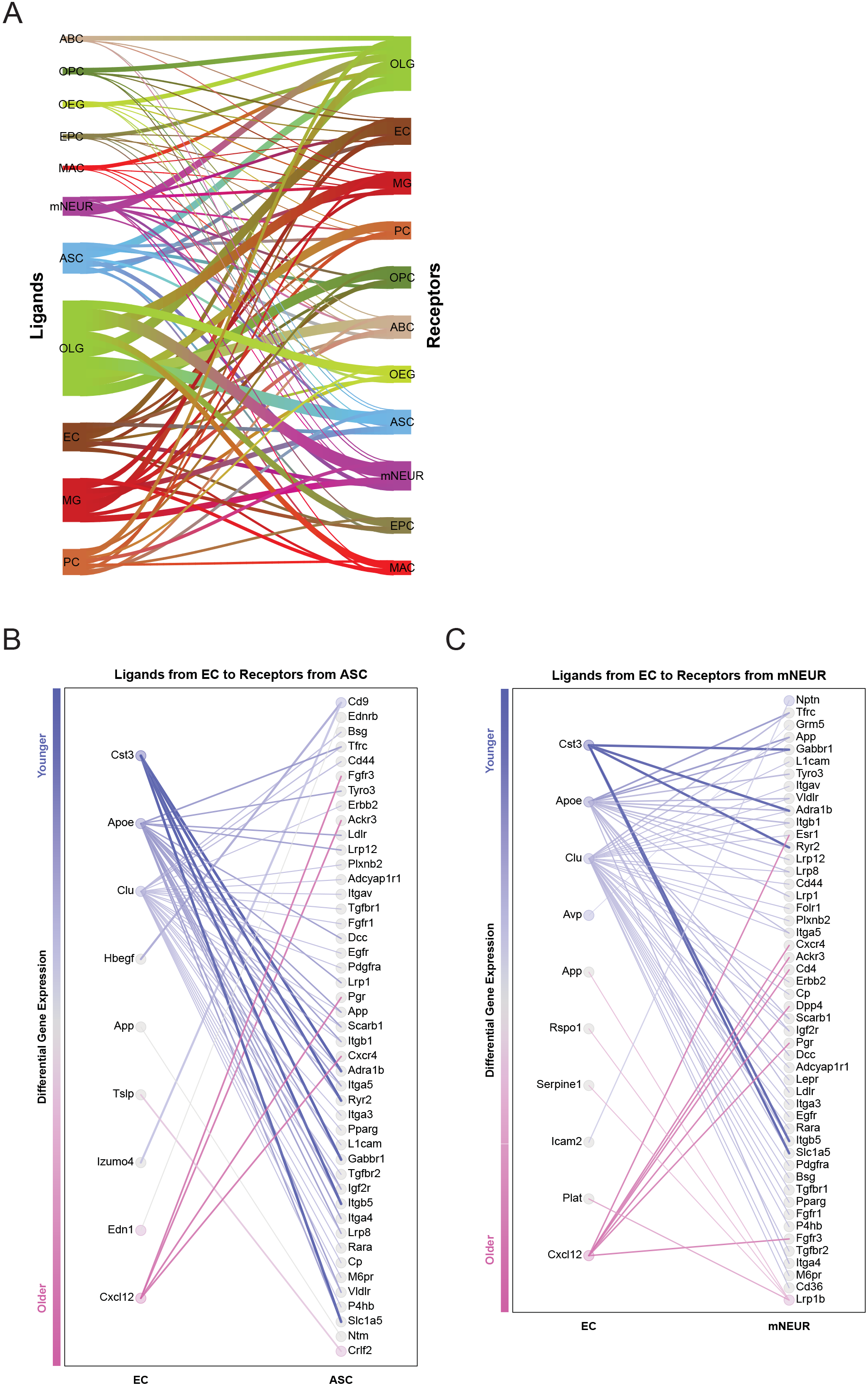
Aging-related changes in intercellular communication. (A) Sankey diagram showing a simplified subset of a network of ligand-receptor interactions among 11 major brain cell types. As a paradigm, the panel in (B) shows aging-related ligands produced and secreted by EC with receptors expressed in ASC, while the panel in (C) shows aging-related ligands produced and secreted by EC with receptors expressed in mNEUR. In both panels, nodes represent ligands or receptors expressed in the denoted cell type, and edges represent protein-protein interactions between them. Node color represents differential gene expression score (signed negative log of FDR) scaled for each cell type, such that the most significantly upregulated genes in aging are magenta, and downregulated are blue. Edge color represents the sum of scores from each contributing node, and only edges with a summed score corresponding to a combined FDR less than 10^−6^ are included in the figure. Edge width and transparency represents the change in each ligand-receptor interaction with aging, calculated by summing the differential expression scores of the participating ligand and receptor for each edge (see Supplementary Methods).

Among the examined cell populations, we highlight the ligand-receptor changes in EC, not only because they exhibited numerous aging-related changes, as mentioned above (see Fig. 2B-C and Fig. S15A), but also due to their unique ability to interact directly with factors from both the brain and the periphery. Network analysis showed that both cystatin C (*Cst3*; an aging-downregulated gene), and stromal cell-derived factor 1 (*Cxcl12*; an aging-upregulated gene) (see Fig. 3B), which have been previously linked to multiple pathologies ^40–43^, are common mediators of crosstalk between vascular cells and many brain cell types (Fig. 6B-C and Table S9). This signifies that their aging-related changes may modulate, either synergistically or separately, important yet unidentified aging-related processes in the brain parenchyma.

## DISCUSSION

The transcriptomic database reported here is the first to examine the aging process in the mammalian brain at a single-cell level. In this study, we first investigated the cellular complexity of the mouse brain and showed that cell identity and composition is overall maintained with aging. More specifically, we found that the numbers of cells within most of the cell types did not change radically as a fraction of total brain cells with age, which is in line with previous findings ^44–48^. Nonetheless, we did observe the previously reported aging-related decline of OPC ^49^, NRP ^35,36^ and ImmN ^36,50^. Of note, it seems possible that additional work focused on this topic itself will reveal additional population shifts in subtypes of cells, particularly those occurring in specific regions of the brain. We then compared young and old cells and observed a noticeable aging-related cell-to-cell transcriptional variation within certain cell populations. However, our data did not show a universal aging-related change in cell-to-cell transcriptional variability across all cell types. That is, gene transcription, in particular cell populations, does not necessarily become more variable with aging. This is in line with Warren et al. ^51^, but in contrast to other studies that suggested increased transcriptional variability is a common feature of aging ^34,52^.

By aggregating all our sequenced single cells and performing DGE analysis comparable to prior bulk sequencing studies, we validated many of the previously identified aging-related genes ^5,7,37^ and extended the list to include numerous additional gene signatures. We then utilized single cell type DGE analyses to reveal the primary cell type(s) generating these signatures. The fine resolution provided by scRNA-seq further allowed us to detect changes in specific cell populations that would otherwise be masked by bulk sequencing techniques. More specifically, single cell type DGE analyses yielded a large number of aging-related genes that are: (a) commonly regulated among cell types, (b) specific to certain cell types and (c) discordant between cell types. To the best of our knowledge, only a small fraction of the genes reported here have been previously associated with brain aging.

Interestingly, our data analysis revealed different patterns of aging across cell populations. We found that certain aging-related genes and pathways are differentially regulated across cell types. For example, we provide evidence that, with aging, expression of ribosomal genes is regulated in opposite directions between groups of cell types. Data from both DGE and pathway analyses showed that the majority of differentiated brain cell types exhibited an aging-driven upregulation of ribosomal genes, while the majority of progenitor cells exhibited the opposite regulation. This paradoxical bidirectional regulation of ribosomal genes with aging is noteworthy.

Over the past decades, it has been clearly shown that the attenuation of protein synthesis by dietary restriction or genetic manipulation of translation-associated genes, including those encoding ribosomal subunits, increases the lifespan of multiple species ^53^. Notably, the down-regulation of ribosomal genes and bulk protein synthesis has been long considered as a hallmark of aging ^54,55^. It appears that aging-driven down-regulation of ribosomal genes had been widely accepted, mostly based on transcriptomic studies in yeast ^56,57^. However, a number of studies in model organisms and humans have presented conflicting results ^6,58–60^. Zahn et al. reported an aging-driven upregulation of ribosomal genes in human brain and muscle tissues ^37^ and, in a later study, reported an aging-driven upregulation of ribosomal genes in mouse neuronal tissues ^6^ but a downregulation of the same genes in non-neuronal tissues ^6^. Moreover, recently published transcriptomic studies showed an aging-related downregulation of ribosomal genes in both ASC ^14^ and NSC ^61^, and an upregulation in MG of both aged ^62,63^ and diseased brains ^62,64^. Intriguingly, a very recent study reported increased ribosomal biogenesis and activity as hallmarks of premature aging in human fibroblasts ^65^. A possible explanation for this is that cells with different metabolic expenses are affected differently by aging, thus inducing alternative feedback loops to partially compensate for loss of translational efficiency and protein synthesis. Collectively, these data indicate that the aging process may not be identical in all cell types, which is in line with our findings and with a recent transcriptome analysis of the *Drosophila* brain that showed a differential aging trajectory in the transcriptional profile of neurons and glial cells ^59^. In short, it is not yet clear whether the regulation of translation-associated genes is causative of aging or the consequence of physiological changes accompanying aging ^53^, or both depending on species, tissue and cell type. However, our work demonstrating that ribosomal biogenesis is one of the aging-related pathways that is differentially regulated across different cell types may help to reconcile seemingly conflicting studies.

Lastly, we created a roadmap of intercellular communication in the brain by generating detailed information on ligand-receptor interactions that change with aging across nearly all brain cell types. This is also of high importance as recent findings from our lab ^66^ and others ^39,67–69^ have shown that certain secreted factors, either derived from brain parenchyma or blood, are able to modulate brain aging, degeneration and rejuvenation. Thus, the discovery of novel factors, their source, and their targets are emerging areas of importance in the aging field and will be crucial for understanding brain physiology in both health and disease ^3^. We foresee the extension of this network by including data from blood proteomic analyses and transcriptomic data from both mouse disease models and heterochronic parabiosis experiments ^66^, to allow further investigation that may help the identification of novel anti-aging therapeutic targets.

Our findings, in agreement with recent studies, underline the sensitivity and power of single-cell transcriptomics not only to reveal differences in cell identities, but also to reveal changes within individual cell types under different treatments and conditions ^23,26,27,70^, including organismal aging ^59,60^. As single-cell sequencing technologies continue to mature, some of the technical and experimental limitations that we encountered will be improved upon. These include: (a) potential sampling problems resulting from the enzymatic dissociation of the brain that may be overcome using single-nuclei sequencing approaches ^20,71^; (b) potential age-associated biases in response to dissociation, cell encapsulation, and other procedures that might drive transcriptional differences between experimental groups; (c) the relatively shallow depth of sequencing limiting the analysis to highly transcribed genes; (d) the relatively small number of cells sequenced compared to the total size of brain tissue restricting the comparative analyses to more abundant cell populations. Furthermore, our data could not reveal sex-specific gene expression changes as only male mouse brains were analyzed. Nonetheless, our work identified aging-related changes in nearly all mouse brain cell types and revealed different patterns of aging across different cell populations, many of which we validated in this study. Thus, while there may be hallmarks of aging that occur in most cell types, such as mitochondrial dysfunction and loss of proteostasis ^1,4,54^, our data argue against the hypothesis that aging induces a single universal molecular program in all cells and tissues ^16^. Future studies will assist in deciphering the exact molecular mechanisms and dynamics by which particular cell types, tissues and organisms respond to normal aging.

Collectively, as a resource to the aging community, we provide a comprehensive dataset of genes, pathways and ligand-receptor interactions with aging-related variation for all the cell types identified. We expect that, beyond the valuable exploration of aging signatures and novel insights regarding the aging process, our data will be used as a reference for a series of other applications. For example, we showed that numerous putative cell specific markers (like *Gfap*) change with aging. Thus, the purification or investigation of cells based on single marker genes maybe faulty in the context of aging. Similarly, our data revealed that the transcript levels of certain housekeeping genes (like *Gapdh*) change dramatically with aging in many cell types, so they cannot be reliably used for quantitative purposes. Finally, we expect that these data will help to advance a variety of efforts towards understanding and modulating the aging process and exploring molecular and cellular therapeutic targets for aging-related neurodegenerative diseases. Furthermore, we hope that our data will serve as a benchmark from which to carry out future studies with higher spatial and temporal resolutions.

## ACKNOWLEDGEMENTS

We would like to acknowledge Tomotaka Okino and his team at Ono Pharmaceuticals for fruitful discussions and useful suggestions during the progress of this work. We are also grateful to Kathleen Pfaff, Francesca Rapino, Natalia Rodriguez-Muela, Adam Freeman, and Jane LaLonde for their helpful advice in different aspects of our work and/or for reviewing the manuscript. The work was supported by Ono Pharmaceutical Co., Ltd (L.L.R.), the Stanley Center for Psychiatric Research and the Klarman Cell Observatory (J.Z.L). The funders had no role in the study design, experiments performed, data collection, data analysis and interpretation, or preparation of the manuscript.

## AUTHOR CONTRIBUTIONS

M.X., S.L.L., S.M.B. and L.L.R. conceived the study; M.X., S.L.L., S.M.B., J.Z.L. and L.L.R. designed the study; M.X., X.A., D.D. and L.N. performed the single cell RNA-seq experiments; S.L.L., S.K.S. and J.Z.L. processed the single cell RNA-seq data; M.X. and S.L.L. analyzed the single cell RNA-seq data; B.T.I. and G.D.B created the ligand-receptor interactions network; B.A.M. performed the RNAscope ISH experiments; C.O. performed the IHC experiments; M.X. and Z.N. performed the flow cytometry experiments; M.X. performed the qRT-PCR experiments; M.X. and V.L.B. performed the bulk RNA-seq experiments; S.L.L and V.L.B. processed the bulk RNA-seq data; M.X. and S.L.L. analyzed the bulk RNA-seq data; R.I. and G.D.B. provided the cell-cell interaction dataset; S.R.L., A.R., G.D.B., J.Z.L. and L.L.R. supervised the study; J.Z.L. and L.L.R. secured funding; M.X., S.L.L., S.M.B. and L.L.R. wrote the manuscript; All authors reviewed the manuscript and approved its submission.

## METHODS

### Animals

C57BL/6J mice (JAX #000664) were housed in the Harvard Biolabs Animal Facility under standard conditions. All experimental procedures were approved in advance by the Animal Care and Use Committee of Harvard University (AEP #10-23) and are in compliance with federal and state laws. Young male mice were used at 2-3 months of age, and old male mice at 21-22 months of age.

### Brain tissue dissociation

Here we modified existing dissociation protocols and developed a new protocol that enables the isolation of intact living cells from both young and old mouse brains in less than 1 hour. Briefly, mice were CO_2_-anesthetized and then rapidly decapitated. Brains were extracted, and hindbrain regions were removed. Meninges were also removed, and the remaining brain tissue was dissociated into single cells using the Adult Brain Dissociation kit (Miltenyi Biotec #130-107-677) with these modifications: (a) the tissue was manually dissociated following the basic steps of the protocol described in the Neural Tissue Dissociation Kit (Miltenyi Biotec #130-092-628); (b) 5% (w/v) trehalose (Sigma Aldrich #T0167) was added in all buffers to ensure higher cellular viability ^81^; (c) half concentration of papain was used, and the digestion was performed at 33-35°C; (d) the enzymatic reaction was quenched using ovomucoid protease inhibitor, as described in the Papain Dissociation System (Worthington #LK003182); (e) cell clusters were removed by serial filtration through pre-wetted 70um (Falcon #352350) and 40um (Falcon #352340) nylon cell strainers; (f) myelin debris and erythrocyte removal steps were omitted to prevent any bias in the recovered cell yields; (g) all centrifugations were performed at 220xg for 8min at 4°C. After dissociations, cells were kept on ice for no longer than 1 hour until further processing.

### Single-cell RNA-sequencing

For the scRNA-seq experiments, 8 young and 8 old mouse brains were analyzed, with 2 animals sacrificed per day. Brain cells were processed through all steps to generate stable cDNA libraries. Briefly, after dissociation, cells were diluted in ice-cold PBS containing 0.4% BSA at a density of 1,000 cells/ul. For every sample, 17,400 cells were loaded into a Chromium Single Cell 3’ Chip (10x Genomics) and processed following the manufacturer’s instructions. Single-cell RNA-seq libraries were prepared using the Chromium Single Cell 3’ Library & Gel Bead kit v2 and i7 Mutiplex kit (10X Genomics). Libraries were pooled based on their molar concentrations. Pooled libraries were then loaded at 2.07 pM and sequenced on a NextSeq 500 instrument (Illumina) with 26 bases for read1, 57 bases for read2 and 8 bases for Index1. Cell Ranger (version 1.2) (10X Genomics) was used to perform sample de-multiplexing, barcode processing and single cell gene unique molecular identifier (UMI) counting, while a digital expression matrix was obtained for each experiment with default parameters ^82^, mapped to the 10x reference for mm10, version 1.2.0. After the initial sequencing, the samples in each pool were re-pooled based on the actual number of cells detected by Cell Ranger (Fig. S2A), aiming to sequence each sample to a similar depth (number of reads/cell) (median: 40,007; Fig. S2C). Multiple NextSeq runs were conducted to achieve over 70% sequencing saturation as determined again by Cell Ranger (median: 75%; Fig. S2F).

### Raw data processing and quality control for cell inclusion

Basic processing and visualization of the scRNA-seq data were performed using the Seurat package (version 2.3) in R (version 3.3.4) ^83–85^. Our initial dataset contained 50,212 cells with data for 19,607 genes. The average numbers of UMI (nUMI) and non-zero genes (nGene) were 2876.70 and 1112.56 respectively. The data were log normalized and scaled to 10,000 transcripts per cell. Variable genes were identified with the FindVariableGenes() function with the following parameters used to set the minimum and maximum average expression and the minimum dispersion: x.low.cutoff = 0.0125, x.high.cutoff = 3, y.cutoff = 0.5. Next, principal component analysis (PCA) was carried out, and the top 20 principal components (PCs) were stored, which is the default number in Seurat. Clusters were identified with the FindClusters() function using the shared nearest neighbor (SNN) modularity optimization with a clustering resolution set to 1.6. All clusters with only one cell were removed. This method resulted in 40 initial clusters. Data for all cells are provided in Fig. S3A with colors representing each of the clusters. For initial quality control filtering, we selectively removed entire clusters with the majority of cells having greater than 30% mitochondrial RNA, under 1,000 detected transcripts, or under 500 unique genes. Finally, we filtered the remaining individual cells using the following parameters: minimum percent mito = 0, maximum percent mito = 30%, minimum number of UMI = 250 maximum number of UMIs = 6000, minimum number of nGene = 250, and maximum number of nGene = 6000 to exclude outliers. Finally, we removed any genes that were only detected in fewer than 3 cells. After initial quality control (QC), we maintained a total of 38,244 cells and 14,699 genes. Data for all cells are provided in Fig. S3B with black representing excluded cells and grey the included cells. The average nUMI, non-zero genes, percent mitochondrial RNA, and percent ribosomal RNA were 3199.12, 1284.08, 8.33%, and 6.94% respectively. PCA was again carried out, and the top 20 PCs were retained. The clustering was again performed with the clustering resolution now set to 2.0. This method resulted in 55 initial clusters. The final pre-processing stage was to remove likely doublet artifacts arising from the co-capture of multiple cells in one droplet. This step occurred following an initial round of determination of cell type identity as described in the next section. We first searched for the top differential markers for each identified cluster/sub-cluster using the FindMarkers() function (Table S3). Then, we defined doublets/multiplets as any cluster in which >30% of its cells express at least 5 of the top 10 genes specific for the initially identified cell type and any other cell type outside of the class of cell types it is associated with (see below for details on cell type classes). These clusters were removed from downstream analysis and clustering was again performed. Ultimately, we included 37,089 cells representing 38 clusters (Fig. S4).

### Determination of cell type identity

We used multiple cell-specific/enriched gene markers that have been previously described in the literature to assist in determining cell type identity (Fig. 1B-D). We then arranged all the identified cell types based on their expression profile, lineage, function and topology into 6 classes of cells (Fig. S7). For each group we re-clustered the subcategorized cell types following the same strategy (top 20 PCs using a clustering resolution of 2.0). Only for the neuronal lineage, which has an increased complexity in terms of cell subtypes, we utilized the top 40 PCs to yield more separated clusters. The annotation of sub-clusters was performed similarly to identification of the main cell clusters.

### Differential gene expression analysis

After initial quality control pre-processing and determination of cellular identities, we utilized the MAST package (version 1.6.1) ^86^ in R (version 3.3.4) to perform differential gene expression (DGE) analysis. MAST generated p-values, fold changes (FC), and logFC (based on natural log of the fold changes) using a hurdle model with normalized nUMI as a covariate. It is worth mentioning that due to shrinkage in the Bayes approach leveraged by MAST, we were able to detect significance in very small changes in transcription but there was also an underestimation of fold change. This is especially noticeable when comparing fold change between MAST calculations and traditional TPM-based calculations for genes with low expression levels. Additionally, the DGE techniques employed here have more power to assign significance of subtle changes in highly transcribed genes and therefore our results may underrepresent changes in lowly transcribed genes. Finally, our ability to establish a baseline level of transcription is proportional to the number of cells measured and thus more subtle changes in abundant populations can be deemed significant.

### Pathway analysis

Gene set enrichment analysis (GSEA) was performed to identify cellular pathways and processes associated with aging ^38^. Analysis was carried out using the GSEA package (version 3.0) (Broad Institute), following the protocol described in Reimand et al. 2017 ^87^. Briefly, prior to the analysis, genes for every distinct cell population were ranked according to their differential gene expression changes and significance (young vs. old). 2 pre ranked gene lists were generated for each cell type: (1) with all genes transcribed, and (2) without the highly abundant mitochondrially-encoding genes and ribosomal genes. All of these pre-ranked gene lists were then used as an input, while 5 gene datasets [Hallmark pathways; GO biological processes; KEGG; BioCarta; Reactome (versions 6.1)] were used as a reference. One thousand random permutations were performed to calculate the p-values for each pathway. Only gene sets with p<0.05 and q<0.25 were considered as significantly enriched. To overcome redundancy and help interpretation of the analysis, we grouped terms over-representing the same pathway using the Cytoscape software (version 3.5.1) and the AutoAnnotate app (version 1.2) ^88^. Pathways belonging to similar biological processes were also grouped together for easier navigation (Table S7). Unless otherwise stated, expression heatmaps for specific pathways or processes were generated using the raw normalized expression (TPM) values. More specifically, for each value in a row of expression the mean of the row was subtracted followed by division by the row’s standard deviation.

### Network analysis

Cell-cell interactions were predicted by a method similar to that described by Kirouac *et al.* ^89^. First, a cell communication interactome was created, collecting known protein-protein interactions between receptor, ligand, and ECM proteins. Receptor genes were defined based on a set of GO terms (GO: 0043235 - receptor complex; GO: 0008305 - integrin complex; GO: 0072657 - protein localized to membrane; GO: 0043113 - receptor clustering; GO: 0004872 - receptor activity; GO: 0009897 - external side of plasma membrane) and UniProt (search term: "Receptor [KW-0675]" GO: 0005886 organism: human). Ligand genes were defined based on a GO term (GO: 0005102 - receptor binding) and the set of proteins labeled as secreted in the Secretome dataset (https://www.proteinatlas.org/humanproteome/secretome) ^90^. ECM genes were defined based on a set of GO terms (GO: 0031012 - extracellular matrix; GO: 0005578 - proteinacious extracellular matrix; GO: 0005201 - extracellular matrix structural constituent; GO: 1990430 - extracellular matrix protein binding; and GO: 0035426 - extracellular matrix cell signalling). Gene lists were manually curated to correct or remove genes that were misclassified. Using the curated list of receptors, ligands, and ECM genes, known protein-protein interactions were collected from iRefindex (version 14) ^91^. Pathway Commons (version 8) ^92^, and BioGRID (version 3.4.147) ^93^, keeping only those occurring between genes from the different classes (ligand, receptor, ECM). This dataset is available at https://baderlab.org/CellCellInteractions. To predict cell-cell interactions, the ligand-receptor interaction set was filtered for genes detected to be expressed at the mRNA transcript level in our cell types. To investigate aging-related perturbations in these putative cell-cell interaction networks, differential gene expression scores (signed negative log of FDR from the MAST analysis outlined above) were used to build subnetworks for each set of interactions between cell types. In these networks, nodes represent ligands or receptors expressed in the denoted cell type, and edges represent protein-protein interactions between them. Edges were included if the sum of scores from each contributing node represented a combined FDR of less than 10^−6^. Nodes were colored to represent the differential gene expression scores scaled for each cell type. Edge color represents the sum of scores from each contributing node. Edge width and transparency represents the magnitude of the summed scores from each contributing node, scaled for each comparison between cell types.

### Flow cytometry analysis

For the simultaneous isolation and purification of ASC, EC, and MG, we developed a multicolor flow cytometry approach. Briefly, dissociated cells from each brain were pelleted (220xg, 8min, 4°C) and resuspended in 1ml ice-cold labeling buffer (HBSS without calcium and magnesium, 0.1% BSA, 2mMEDTA, 5% trehalose, 1% GlutaMAX). Cells were incubated with 100ul of FcR blocking reagent (Miltenyi Biotec #130-092-575) for 12min at 4°C under continuous rotation, and then labeled with the following antibodies: APC anti-ACSA-2 (Miltenyi Biotec #130-102-315) for ASC; BV786 anti-CD31 (BD Biosciences #740870 and BD Biosciences #740879) for EC; and BV510 anti-CD11b (BD Biosciences #562950) for MG. Cells were also incubated with the following antibodies targeting unwanted cell populations: PE anti-CD200 (BioLegend #123808) for mNEUR; Alexa Fluor 488 anti-04 (R&D Systems #FAB1326G) for OLG; and BV605 anti-CD140a (BD Biosciences #740380) for OPC. This step is critical as it helps to exclude these cells during sorting, thus minimizing cross contamination events. After 12min of incubation at 4°C (in dark conditions), cells were washed extensively, pelleted and resuspended in ice-cold FACS buffer (HBSS containing calcium and magnesium, 0.5% BSA, 5% trehalose, 1% Glutamax) in a volume of 25ml per brain (5 FACS tubes). To exclude cellular debris and dead cells, 15min before sorting, 10uM Calcein Blue AM (BD Biosciences #564060) was added to the FACS tubes to stain live cells. Calcein+ cells were then sorted using a Moflo Astrios instrument (Beckman Coulter) with a 70um nozzle at 60psi. Gates were set manually by using compensation beads (Life Technologies #A10497) and appropriate control samples, and data were analyzed with FlowJo software (version 10). To minimize RNA degradation, cells were collected directly in RL buffer (Norgen Biotek #48500) supplemented with 10% BME, in a 1:1 final ratio (50% lysis buffer : 50% cells in sheath fluid; the PBS-based solution that is derived from the flow cytometer). After sorting, cell lysates were snap frozen and stored at −80°C for up to 1 month until further processing.

### RNA extraction

Total RNA was extracted from sorted cells using the total RNA purification plus Micro kit (Norgen Biotek #48500) following the manufacturer’s instructions. Prior to RNA extraction, a chloroform extraction step was included to remove myelin debris/lipids, as well as, an on-column DNase digestion step (Qiagen #79254) to remove genomic and mitochondrial DNA. For all samples, RNA concentration was determined using a Qubit Fluorometer (Invitrogen), while RNA purity and integrity were evaluated with a BioAnalyzer instrument (Agilent). After extraction, RNA was immediately stored at −80°C for no longer than a month until further processing.

### Bulk RNA sequencing

For the bulk RNA-seq experiments, sorted/purified cells from 8 mouse brains (4 young and 4 old) were analyzed. Bulk RNA-seq was performed using a modified version of the SCRB-Seq that was originally developed for single cell RNA-seq analysis ^94^. Briefly, polyadenylated RNA, from total RNA (7.5-25ng; RIN values >6.5) extracted from our FACS-purified cells, with ERCC Spike-in control Mix A (Ambion) at 10^−6^ final dilution, were converted to cDNA and decorated with universal adapters, sample-specific barcodes and UMI using a template-switching reverse transcriptase. Decorated cDNA was then pooled, amplified and prepared for multiplexed sequencing (NextSeq500, Illumina) using a modified transposon-based fragmentation approach that enriched for 3’ ends and preserved strand information.

### Bulk sequencing data analysis

Post-sequencing quality control on each of the libraries was performed to assess coverage depth, enrichment for messenger RNA (exon/intron and exon/intergenic density ratios), fraction of rRNA reads and number of detected genes using bespoke scripts. Second sequence reads were aligned against the murine genome mm9 using bwa mem (version 0.7.10-r789) (http://bio-bwa.sourceforge.net/). Gene expression was estimated based on reads mapping near the 3’ end of transcripts using ESAT ^95^, based on the mm9 Refseq annotation, with flags java -Xmx128G -task score3p -wLen 50 -wExt 5000 -wOlap 0 -sigTest 0.01 -multimap ignore. Results were summarized as counts per million mapped reads (CPMs), merged across samples, log-transformed and subjected to hierarchical clustering and visualization. For ERCC quantification, reads were mapped against the ERCC sequences using STAR (version 2.5.1b) ^96^ with flags --runMode alignReads --runThreadN 8–outSAMtype BAM SortedByCoordinate–outFilterType BySJout --outFilterMultimapNmax 20 -- outFilterMismatchNmax 999 --alignIntronMin 10-alignIntronMax 1000000 --alignMatesGapMax 1000000 --alignSJoverhangMin 8 --alignSJDBoverhangMin 1 --quantMode TranscriptomeSAM. Bam files from STAR were sorted and indexed with samtools ^97^ and counts were retrieved from the indices using idxstats. Differential gene expression (DGE) analysis ^98^ was performed in R (version 3.2.3) using Bioconductor’s DESeq2 package (version 3.7) ^99^. Dataset parameters were estimated using the estimateSizeFactors(), and estimateDispersions() functions; read counts across conditions were modeled based on a negative binomial distribution and a Wald test was used to test for differential expression (nbinomWaldtest(), all packaged into the DESeq() function), using the age as a contrast.

### Quantitative Real time PCR

For the quantitative Real-Time PCR (qRT-PCR) experiments, sorted/purified cells from 11 mouse brains (5 young and 6 old) were analyzed. Briefly, RNA samples with RIN values >6.5 were reverse transcribed into cDNA using the iScript cDNA synthesis kit (Bio-Rad #170-8891) following the manufacturer’s instructions. The resulting cDNA was then processed for qRT-PCR analysis using pre-designed primers (Integrated DNA Technologies) (Table S10) and the Fast SYBR Green Master Mix (Life Technologies #4385614) in a QuantStudio 12K Flex Real-Time PCR System (Applied Biosystems). Before data analysis, we examined the melting curves for each reaction and included only those with a single peak at the expected melting temperature. The fold-change (FC) in gene expression was determined by the 2^−DDC^_T_ method ^100^, and all values were normalized to the endogenous expression of *Vcp*; a housekeeping gene that has been proposed for calibration in quantitative experiments ^101^. Our analysis showed that in the vast majority of cell populations its levels remain unaltered with aging, in contrast to other more commonly used genes, like *Gapdh*. Samples with *Vcp* Ct values >28.5 were excluded from our analysis. Each sample was repeated in technical duplicates on 3-6 biological replicates.

### RNAscope In Situ Hybridization

RNAscope fluorescent in situ hybridization was performed on fresh-frozen brain tissue from 6-8 mice (3-4 young and 3-4 old). For sample preparation, mice were CO_2_-anesthetized, and brains were rapidly extracted and embedded in OCT (Tissue Tek) on dry ice, and then stored at −80°C until further processing. We collected 14μm cryostat sections and RNAscope hybridizations were carried out according to the manufacturer’s instructions, using the RNAscope Multiplex Fluorescent Manual Assay kit (Advanced Cell Diagnostics). Briefly, thawed sections were dehydrated in sequential incubations with ethanol, followed by 30 min Protease IV treatment and washing in 1x PBS. Appropriate combinations of hybridization probes were incubated for 2 hours at 40°C, followed by four amplification steps, DAPI counterstaining, and mounting with Prolong Gold mounting medium (Thermo Fisher Scientific #P36930). Probes against *Rpl6, Malat1*, and *Meg3* (Advanced Cell Diagnostics) were tested. For each mouse, 3-4 bregma-matched sections were imaged. Images were acquired using a Zeiss LSM 880 Confocal Microscope and represented as maximum intensity projections of acquired confocal z stacks. Analysis was done using the CellProfiler software (version 3) ^102^. Only puncta with a diameter between 4-8 pixel-units that were located within the nuclei (for Malat1 and Meg3) and perinuclear space (for Rpl6) were quantified.

### Immunohistochemistry

For immunohistochemistry experiments, 8 mouse brains (4 young and 4 old) were processed. Mice were perfused intracardially with 1x PBS followed by 4% paraformaldehyde (PFA). Brains were removed and embedded in 3% agarose. 30μm-thick free-floating coronal sections were cut in a vibrating microtome and were kept in 1x PBS with 0.1% sodium azide at 4°C until staining. Sections were washed thoroughly in 1x PBS and incubated in a permeabilization/blocking solution (10% normal goat or donkey serum, 0.2% Triton X-100, 1x PBS) for 1 hour at room temperature. Sections were incubated overnight at 4°C with the following primary antibodies in blocking solution: goat polyclonal anti-SPARC (R&D Systems #AF942), rabbit polyclonal anti-IBA1 (Wako #019-19741), and goat polyclonal anti-IL33 (R&D Systems#AF3626). Alexa Fluor secondary antibodies (Invitrogen) were then used for detection of primary antibodies in 1% normal goat or donkey serum, 1x PBS for 1-2 hours at room temperature. Hoechst 33342 was used to label nuclei. Imaging was performed using Zeiss ELYRA super-resolution confocal microscope at 20x and 40x magnifications. Images were visualized using Zeiss Zen software. For each mouse, 3-4 bregma-matched sections were imaged. Images were represented as maximum intensity projections of acquired confocal z-stacks. Analysis was done using Image J software (version 1.49)

### Statistical analysis

All statistical analyses were performed using R (version 3.3.4) or GraphPad Prism (version 7.04). Unless otherwise stated, to generate p-values for cell counts and other metrics/variables (nGene, nUMI, CV) we used the Mann-Whitney *U* test ^103^. All p-values modified to a false discovery rate (FDR) of 5% using the Benjamin-Hochberg precedure ^104^.

### Data availability

To facilitate the exploration and utilization of our processed data, they will be publicly available through the Broad Single Cell Data Portal using this link: https://portals.broadinstitute.org/singlecell/study/aging-mouse-brain

